# DNA Methylation Explains a Subset of Placental Gene Expression Differences Based on Ancestry and Altitude

**DOI:** 10.1101/261974

**Authors:** William E. Gundling, Priyadarshini Pantham, Nicholas P Illsley, Lourdes Echalar, Stacy Zamudio, Derek E. Wildman

## Abstract

**Objectives**: The most pronounced effect of high altitude (>2700m) on reproductive outcomes is reduced birth weight. Indigenous Bolivians (Andean Native Americans) residing for generations at high altitudes have higher birth weights relative to more recent migrants of primarily European ancestry. Previous research demonstrated that the placenta is a key contributor to the preservation of Andean birth weight at high altitude. Our current research investigated how gene expression and epigenetics contributes to the conservation of birth weight at high altitude by examining mRNA expression and DNA methylation differences between placentas of Andeans and those of European ancestry residing at high and low altitude.

**Methods**: Genome-wide mRNA expression and DNA methylation of villous placenta tissue was quantified utilizing microarray technology. Subjects were of Andean and European ancestry and resident at high (3600m) or low (400m) altitudes. Differentially expressed genes (DEGs) associated with altitude or ancestry were identified (FDR<0.1, |fold change|>1.25). To predict which DEGs could be regulated by methylation we tested for correlation between gene expression and methylation values.

**Results**: 69 DEGs associated with altitude (n=36) or ancestry (n=34) were identified. Altitude-associated DEGs included members of the AP-1 transcription factor family. Ancestry-associated DEGs were implicated in inflammatory pathways and associated with pro-angiogenic macrophages. More ancestry-associated DEGs correlated significantly (n=17) (FDR<0.1) with promoter or gene body methylation (p=0.0242) when compared to altitude associated DEGs (n=8).

**Conclusions:** Compared to altitude-associated DEGs, methylation regulates more ancestry-associated DEGs, potentially allowing for rapid modification in the expression of inflammatory genes to attract pro-angiogenic macrophages as a means of promoting placental capillary growth in Andeans, regardless of altitude.

## 1. Introduction

The placenta mediates fetal growth by orchestrating nutrient and gas exchange between the mother and the fetus [1]. We have shown there is altered substrate delivery from placenta to fetus under conditions of chronic maternal hypoxemia (lowered arterial oxygen tension due to high-alitude residence) [10]. High altitude environments contributes to reduced birth weights in humans, with a gradient of reduction in birth weight that corresponds to the evolutionary duration of population exposure to high altitude [2-4]. At ≥3000 meters, Tibetans and Andeans have higher birth weights than more recent migrants such as European and Han (Ethnic Chinese) [3]. The Andeans, with a population history of ~10,000 years at high altitude have birth weights >200 grams higher than Europeans at ≥3000 meters [2, 3, 5]. This suggests a relatively rapid adaptation to the high altitude environment in terms of the promotion of favorable reproductive outcomes among Andeans. Recent analyses have shown Tibetan and Andean populations have undergone natural selection to favor variants of genes associated with the hypoxia response, oxidative metabolism, and possibly regulation of vascular tone [4]. However, the definitive link between gene variants and altered physiological function has only been inferred, as genetic analyses of high altitude populations have relied upon DNA from leukocytes rather than a reproductive tissue such as the placenta.

Reduced birth weight in a population subjected to environmental stress is evidence of negative natural selection because it is associated with increased fetal/neonatal morbidity and mortality [6]. Lowered birth weight is also associated with an increased risk for chronic diseases later in life, including cardiovascular and metabolic diseases [7]. Hence placental adaptations that protect against growth restriction not only benefit offspring survival, but also population-wide health in later life. The accumulated data suggests that at high alitutde there is diminished vascular elasticity, consistent with the reduced uterine arterial and fetal umbilical venous blood flow we and others have reported in high-altitude residents [8-11].

The greater fetal growth in Andean Native Americans is associated with higher uterine arterial and umbilical venous blood flows compared to more recent migrants to high altitude [8, 9]. Our prior work found no relationship between oxygen-associated variables such as maternal and fetal Po_2_, arterial oxygen content and birth outcomes. Instead, placental weight was the strongest correlate of fetal size at birth, accounting for 35% of the variation in birth weight in both Andean and European women at high altitude [9]. Moreover, for any given placental weight, Andean neonates were larger than European, indicative of greater placental efficiency among Andeans. For these reasons we proposed that the ancestry differences we have reported are due to changes in placental function [9, 12]. This was the stimulus for the present study, in which we hypothesize that differences in placental gene expression contribute to the greater growth of the Andean fetus at high altitude.

To our knowledge, global gene expression and DNA methylation in placental tissue has never been examined relative to ancestry, nor to an environmental challenge such as high vs. low altitude. Our approach was to compare the gene expression signatures of placentas from indigenous Andeans to those of more recent migrants, most of whom are the descendants of European colonists residing at high and low altitude in Bolivia (<500 years). The ancestry-associated DEGs may provide insight into how the Andean placenta has adapted to protect the fetus from altitude-associated reduction in birth weight. Altitude-associated DEGs may provide new biomarkers for, or even point towards therapeutic options for hypoxia-related placental pathologies such as preeclampsia and idiopathic intrauterine growth restriction.

We also investigated how DNA methylation correlates with gene expression in the altitude or ancestry-associated DEGs. We reasoned that the ~10,000 year exposure of Andeans to the high altitude environment is not generally considered long enough for evolution by random mutation and subsequent selection. The Andean plateau, while long, is narrow, and undoubetdly prehistoric populations moved between altitudes. DNA methylation may explain how gene expression has been altered relatively rapidly. An environmentally induced methylation event circumvents any need for germ cell mutations and thus would be less “costly” in reproductive terms.

## 2. Methods

### 2.1. Subjects

The data reported here are derived from a subset of pregnancies studied in La Paz, Bolivia (altitude ~3600 m) and Santa Cruz, Bolivia (altitude ~400 m). The original studies were designed to examine maternal and fetal blood flow, oxygen delivery, and consumption in a four-way cross-sectional design comparing indigenous Andeans versus European migrants residing at high and low altitude [8]. All women gave written informed consent to protocols approved by the Bolivian National Bioethics Committee, the San Andreas Mayor University, Instituto Boliviano de Biologia de Altura Consejo Tecnico, and the USA Institutional IRBs.

Women completed an interview and questionnaire in which they identified the birthplace and residence history of themselves, their parents, their grandparents, and those of the baby’s father and his recent ancestors. Three generation residential history in the Altiplano region, Andean surnames, self-identification as, and fluency in Aymara or Quechua initially defined our Andean samples from both high and low altitude. Women who self-identified as Spanish or European, reported no known Andean ancestry, and who documented three generations of low altitude residence were provisionally assigned to the European group from both high and low altitudes. Biogeographic ancestry was evaluated using 133 Ancestry Informative Markers (AIMs) to confirm self-identified ancestry and quantify admixture [9, 13, 14]. The Andean women in this study had biogeographic ancestry profiles that were on average 86% Native American (95% Confidence Interval (CI) 76-92%), and 9% European (95% CI 6-20%). The European sample population had an average of 63% European ancestry (95% CI 54-68%) and 31% Native American (non-Andean) ancestry (95% CI 22-43%). Low levels of sub-Saharan African (2-6%) and East Asian admixture (3-5%) were detected in both ancestry groups. There was no significant difference between altitudes in the measures of geographic ancestry, i.e. Andean individuals at low vs. high altitude did not differ in their proportional admixture, nor did those of European ancestry between the low and high altitude sites. Maternal inclusion criteria were conception, gestation and birth at altitude of residence, singleton pregnancy, a healthy mother age 18-45 with no known chronic disease, a normal glucose tolerance test, and the absence of any known pregnancy complications.

#### 2.1.2. Placental sampling/morphometrics

All placentas were obtained from elective, non-laboring term cesarean deliveries without supplemental oxygen. Placentas were obtained immediately after delivery. The sampling strategy was designed to ensure multiple portions of the placenta were sampled, but also that RNA was collected as quickly as possible. Therefore, a semi-random sampling strategy was used, in which the placenta was visualized as roughly circular and divided into four quadrants. One full-depth sample from approximately the center of each quadrant was excised, and the basal and chorionic plates were removed, and the core villous tissue minced. Tissue from all four quadrants was thoroughly mixed and 200 mg aliquots were flash frozen in liquid nitrogen for later RNA/DNA isolation. Each RNA/DNA aliquot, therefore, contained tissue from four regions of the placenta and was flash frozen in liquid nitrogen less than 30 minutes following delivery. For placental morphometric studies, eight full depth sections were collected (two from each quadrant), formalin fixed and paraffin-embedded. Five of these samples were used for morphometric analyses following the methods originally described by Weibel et al [15, 16], and as applied by Jackson and Mayhew [17-19] and Burton et. al.[20].

Morphometric studies were overseen by DCM in consultation with Dr. Graham Burton of Cambridge University. The fetal membranes and umbilical cord were trimmed and the placentas were weighed. Placental volume was measured by water displacement, and correction for tissue shrinkage was based on maternal erythrocyte diameters [21]. While complete morphometric analyses were undertaken of 20-30 placentas from each of the 4 low and high altitude groups, only data pertaining to vascular variables is considered here in relation to our microarray results.

### 2.2. Nucleotide Isolation and Expression Microarray

RNA was isolated from the core villous tissue of 45 placentas (n=10 Andeans from high altitude, n=12 Andeans born at high altitude but who migrated to low altitude, n=11 European individuals born at low altitude but who migrated to high altitude, and n=12 Europeans individuals born and residing at low altitude). The concentration and optical density of each RNA sample was measured using a Nanodrop spectrometer (Thermo-Fisher, Waltham MA). The quality of the RNA was further examined by calculating the RNA integrity number (RIN) using a Bioanalyzer (Agilent, Santa Clara CA). RIN score for all samples used for microarray array analysis ranged from 7.3 to 4.6. One μg of RNA per sample was supplied to the Applied Genomics Technology Center at Wayne State University (Detroit, MI). The RNA was converted to cDNA and then hybridized to Illumina HumanHT-12v4 Expression BeadChips (Illumina, San Diego, CA) containing 47,231 probe sequences with each probe sequence replicated on 20-30 beads using Illumina’s standard protocol. The average fluorescence intensity value for each probe set was calculated using the Illumina GenomeStudio software (Illumina, San Diego, CA). The probe sequences target more than 31,000 annotated genes. Samples were plated in random order across the different microarray chips on the same day to minimize batch effects. Intensity values for annotated genes were log_2_ transformed and quantile normalized using the Preprocesscore package from Bioconductor software collection in R [22]. Probes located on the X- and Y-chromosomes were excluded because of sex-based dosage-dependent effects on gene expression. We calculated the variance of each of the remaining probes and sorted them from highest to lowest variance and used the top 5% most variable probes for further analysis [23]. All raw files, non-normalized intensity values as well as quantile normalized and log-transformed files have been uploaded to the GEO database (GSE100988).

#### 2.2.1 Determination of Differentially Expression Genes

We tested for differential expression using a two-way analysis of covariance (ANCOVA) with the model, probe expression in proportion to altitude + fetal sex + gestational age or probe expression in proportion to ancestry + fetal sex + gestational age (Supplementary Table 1). P-values were generated from each independent variable and the false discovery rate (FDR) was estimated using the Benjamini-Hochberg (BH) method to adjust for multiple hypothesis testing [24]. A probe sequence was considered to be significantly differentially expressed if it had an FDR<0.1 and an absolute fold change≥1.25.

#### 2.2.2. Validation of Microarray Data

We conducted qRT-PCR using Taqman gene expression assays (Invitrogen, Carlsbad, CA) to validate the microarray results. 500ng of RNA from each sample was reverse transcribed to cDNA using the Superscript III First-Strand Synthesis system (Invitrogen, Carlsbad, CA). qRT-PCR was performed on the StepOnePlus real-time PCR machine (Applied Biosystems, Foster City, CA) with Taqman Fast Advanced Master Mix (Invitrogen, Carlsbad, CA) and fast cycling conditions. We tested six genes that were differentially expressed due altitude or ancestry with qRT-PCR: lymphatic vessel endothelial hyaluronan receptor 1 *(LYVE1*, Taqman probe Hs00272659_m1), stanniocalcin 1 *(STC1*, Taqman probe Hs00292993_m1), fos proto-oncogene *(FOS*, Taqman probe Hs99999140_m1), arachidonate 5-lipoxygenase-activating protein *(ALOX5AP*, Taqman probe Hs00233463_m1), nicotinate phosphoribosyltransferase *(NAPRT*, Taqman probe Hs00292993_m1), and phospholipase A2, group 2A *(PLA2G2A*, Taqman probe Hs00179898_m1). For each gene tested, samples were randomized and run in triplicate, incorporating a multiplexed 18S ribosomal RNA endogenous control *(18S*, Taqman probe Hs99999901_s1) in each well. The Ct values for each sample and gene were analyzed using the 2^-∆∆Ct^ method [25]. The qRT-PCR intensity value (Ct) for each gene was correlated with the intensity value yielded by the gene expression microarray to validate results from the gene expression microarray.

### 2.3. Pathway Analyses

KEGG pathway enrichment analyses were performed separately on the altitude- and ancestry-associated DEGs using the GOstats package provided by Bioconductor [26]. We used a hypergeometric test to determine which pathways were significantly over-represented within our lists of altitude- and ancestry-associated DEGs. A pathway was considered to be significantly over-represented if it had an FDR<0.05.

### 2.4. Methylation Detection and Quantification

We isolated DNA from the same villous placental tissues of the 45 individuals analyzed for gene expression. The DNA samples were bisulfite converted using the EZ DNA methylation kit protocol (Zymo Research, Irvine, CA) and hybridized to the Illumina HM 450k methylation microarray (Illumina, San Diego, CA) and scanned using the Illumina iScan Array Scanning System (Illumina, San Diego, CA) at the Applied Genomic Technology Center at Wayne State University (Detroit, MI). Fluorescence intensity values (i.e. β values) were calculated using the Illumina GenomeStudio software (Illumina, San Diego, CA), and a final report containing information for each probe was generated and preprocessed with the IMA (Illumina Methylation Analyzer) R package [27]. After preprocessing, we used beta-mixture quantile normalization (BMIQ) to normalize type 1 and type 2 probe intensities [28]. β values were logit transformed and quantile-normalized into M values to reduce the heteroscedasticity of the distribution across our samples [29]. To restrict our analysis to sites that could potentially play a regulatory role in differential gene expression, we only analyzed probes located in the promoter or gene body of altitude- or ancestry-associated DEGs. We defined the promoter region as the area located within 1.5kb upstream of the transcription start site (TSS) through the 5’ untranslated region (UTR). We considered probes to be in the gene body if they were located downstream of the 5’UTR and upstream of the 3’UTR.

### 2.5. Correlation between Gene Expression and Methylation

Pearson’s correlation test was used for evaluation of correlation between the log_2_ intensity RNA abundance from gene expression microarrays and the M values of individual methylation probes within the promoter or gene body region of the identified DEGs in samples for which we had both gene expression and DNA methylation data. A correlation between gene expression intensity and M-value was considered to be statistically significant if the Pearson correlation test had an FDR<0.1. FDR was estimated for ancestry- and altitude-associated DEGs separately, similar to how it was estimated in the 2-way ANCOVA for gene expression.

## 3. Results

### 3.1. Subjects / placental morphometrics

The mothers in this study were similar to those of the larger parent study. Andean women were older and had higher gravidity and parity (Table 1). Non-pregnant Body Mass Index (BMI) was similar across all four groups, as was gestational age at delivery, placental weight and volume. Andean women had heavier birth weights than European women, and less reduction in birth weight at high altitude. Because birth weights were lower, despite similar placental size, altitude reduced placental efficiency, reflected in the placenta:birth weight ratio. Morphometric analyses of the placentas revealed greater vascularization in Andean vs. European placentas at high altitude (Table 1). Of the 45 placentas included in our microarray and methylation experiments, 42 were represented in the 105 that were analyzed for morphometrics. Relative to their low-altitude counterparts Andean subjects had an altitude-associated increase in capillary surface area, length and volume of 32%, 28% and 75% respectively. These changes are greater than those in the total sample of low and high-altitude Andean placentae, in which these differences were 25%, 15% and 38% respectively. Among Europeans there was a 0%, 9% and 4% increase in capillary surface area, length and volume respectively. Again, these data differ somewhat from the larger sample of European placentas analyzed (6%, <1%, 23%). However, it must be remembered that the subset of subjects chosen for the genetic analyses represented the highest degree of Native American ancestry among Andeans and of European ancestry among the subjects of low-altitude origin.

**Table 1:** Clinical data for placenta samples. A 2-way ANOVA was used with altitude and ancestry as the independent variables. An interaction term with P < 0.05 indicates that the phenotype associated with the combination of altitude and ancestry differs significantly btween biogeographic ancestry groups. For example, the decrement in birth weight due to altitude is less among Andeans than Europeans, hence the interaction term is significant.

|  | High<br>Altitude<br>Andean | High<br>Altitude<br>European | Low<br>Altitude<br>Andean | Low<br>Altitude<br>European | Altitude<br>p-value | Ancestry<br>p-value | Interaction<br>p-value |
| --- | --- | --- | --- | --- | --- | --- | --- |
| N | 10 | 11 | 12 | 12 |  |  |  |
| Maternal age | 32.6±5.2 | 30.8±3.1 | 29.3±5.4 | 25.9±5.9 | 0.006 | 0.046 | 0.802 |
| Pre-pregnant<br>BMI | 23.9±2.9 | 23.9±3.0 | 26.2±3.5 | 22.8±4.3 | 0.953 | 0.032 | 0.437 |
| Gravidity | 4.4±3.7 | 1.6±0.9 | 3.8±2.5 | 2.3±1.3 | 0.900 | 0.004 | 0.400 |
| Parity | 1.9±1.7 | 0.5±0.5 | 2.3±2.5 | 1.0±1.0 | 0.330 | 0.009+ | 0.843 |
| Neonatal Sex | M=6, F=4 | M=4, F=7 | M=7, F=5 | M= 5, F= 7 |  |  |  |
| Birth Weight(g) | 3444±426 | 3026±272 | 3766±221 | 3621±375 | <0.001 | 0.009 | 0.174 |
| Gestational Age (wks) | 38.7±1.4 | 38.3±1.1 | 39.0±0.7 | 38.4±0.9 | 0.585 | 0.094 | 0.838 |
| Placental Weight (g) | 549±102 | 500±87 | 493±82 | 476±84 | 0.146 | 0.240 | 0.552 |
| Placental volume (cm<sup>3</sup>) | 556±115 | 500±87 | 463±97 | 446±76 | 0.015 | 0.219 | 0.497 |
| Birth to Placental weight ratio | 6.4±0.7 | 6.2±0.9 | 7.8±1.3 | 7.9±1.1 | <0.001 | 0.691 | 0.797 |
| Capillary Length (Km) | 564±245 | 441±161 | 506±142 | 454±146 | 0.654 | 0.336 | 0.139 |
| Capillary surface area (m<sup>2</sup>) | 16.7±4.3 | 12.2±2.6 | 13.5±3.3 | 13.2±3.8 | 0.302 | 0.053 | 0.068 |
| Capillary volume (cm<sup>3</sup>) | 53.1±15.4 | 33.1±10.8 | 37.8±7.9 | 31.0±11.2 | 0.003 | 0.006 | 0.016 |

### 3.2 Placental Gene Expression

In total, we identified 69 DEGs. Thirty-four genes (42 probe sequences) were differentially expressed between individuals of Andean and European descent (Figure 2, Table 2) and 36 genes (36 probe sequences) were differentially expressed between individuals residing at high and low altitude (Figure 2, Table 3). One gene, Carboxypeptidase, Vitellogenic Like *(CPVL)* was differentially expressed assoicated with both ancestry and altitude. There were no significant differences between fetal sexes and across gestational ages among the genes used to test for differentental expression (Supplmentary Table 1). We tested six DEGs *(LYVE1, STC1, FOS, ALOX5AP, NAPRT1*, and *PLA2G2A)* using qRT-PCR and all showed a significant correlation with microarray expression, therefore validating the microarray analysis (Supplementary Table 2). Of the 34 ancestry-associated DEGs, ten had higher expression in European versus Andean placentas and 24 had higher expression in the Andean placentas (Table 2, Figure 2). The ancestry-associated DEGs with higher expression in Andeans included genes in the gene ontology categories inflammatory response (GO:0006954), response to stress (GO:0006950), and the immune response (GO:0006955) such as chemokines c-c motif chemokine ligand 2 *(CCL2)* and c-c motif chemokine ligand 3 like 3 *(CCL3L3)* major histocompatibility complex class I A *(HLA-A)* and macrophage markers CD163 molecule *(CD163)* and lymphatic vessel endothelial hyaluronan receptor 1 *(LYVE1)* [30].

**Table 2:** Fold change of Illumina HT12-v4 gene expression microarray probes that are significantly differentially expressed (FDR > 0.1) as a function of ancestry

| Probe ID | Gene Symbol | Fold Change | FDR |
| --- | --- | --- | --- |
| ILMN_1710752 | *NAPRT* | -2.03 | 0.011 |
| ILMN_1740586 | *PLA2G2A* | -1.7 | 0.072 |
| ILMN_2173294 | *THNSL2* | -1.65 | 0.001 |
| ILMN_3248833 | *RPS26* | -1.64 | 0.001 |
| ILMN_3299955 | *RPS26* | -1.56 | 0.001 |
| ILMN_1739605 | *LYPD3* | -1.56 | 0.001 |
| ILMN_2209027 | *RPS26* | -1.54 | 0.001 |
| ILMN_2310703 | *RPS26* | -1.52 | 0.004 |
| ILMN_1715169 | *HLA-DRB1* | -1.48 | 0.092 |
| ILMN_1685580 | *CBLB* | -1.47 | 0.011 |
| ILMN_2411282 | *QSOX1* | -1.42 | 0.011 |
| ILMN_1687035 | *ADAMTSL4* | -1.32 | 0.035 |
| ILMN_1762294 | *ADAMTSL4* | -1.3 | 0.053 |
| ILMN_1691717 | *RHBDF2* | 1.26 | 0.051 |
| ILMN_1749403 | *TSPAN33* | 1.27 | 0.035 |
| ILMN_1772359 | *LAPTM5* | 1.28 | 0.071 |
| ILMN_1733270 | *CD163* | 1.28 | 0.064 |
| ILMN_1717163 | *F13A1* | 1.29 | 0.094 |
| ILMN_1683792 | *LAP3* | 1.29 | 0.054 |
| ILMN_1721035 | *MS4A6A* | 1.31 | 0.094 |
| ILMN_1785413 | *ATP2C2* | 1.31 | 0.047 |
| ILMN_2352023 | *DSTYK* | 1.32 | 0.094 |
| ILMN_1657373 | *LEPREL1* | 1.33 | 0.071 |
| ILMN_1808114 | *LYVE1* | 1.34 | 0.099 |
| ILMN_1740024 | *NAALAD2* | 1.35 | 0.051 |
| ILMN_1670490 | *PDPN* | 1.36 | 0.094 |
| ILMN_2203950 | *HLA-A* | 1.38 | 0.094 |
| ILMN_2200659 | *SNHG5* | 1.38 | 0.088 |
| ILMN_1808245 | *SBSPON* | 1.38 | 0.072 |
| ILMN_1662419 | *COX7A1* | 1.39 | 0.072 |
| ILMN_1682928 | *CPVL* | 1.39 | 0.015 |
| ILMN_2379599 | *CD163* | 1.4 | 0.027 |
| ILMN_2130441 | *HLA-H* | 1.4 | 0.086 |
| ILMN_2186806 | *HLA-A* | 1.41 | 0.060 |
| ILMN_1722622 | *CD163* | 1.41 | 0.035 |
| ILMN_2400759 | *CPVL* | 1.42 | 0.011 |
| ILMN_1797875 | *ALOX5AP* | 1.42 | 0.094 |
| ILMN_1758164 | *STC1* | 1.43 | 0.004 |
| ILMN_1772459 | *RPS23* | 1.44 | 0.010 |
| ILMN_1776967 | *DNAAF1* | 1.46 | 0.089 |
| ILMN_2105573 | *CCL3L3* | 1.53 | 0.099 |
| ILMN_1720048 | *CCL2* | 1.56 | 0.035 |

Among the 36 altitude-associated DEGs, 24 DEGs had higher expression levels in placentas of individuals residing at high altitude compared to those residing at low altitude, whereas 12 DEGs were more highly expressed in placentas of individuals residing at low altitude (Table 3, Figure 1). DEGs with decreased expression at high altitude include fos *(FOS)*, fosb *(FOSB)*, and jun *(JUN)* proto-oncogenes, which form part of the activator protein 1 (AP-1) transcription factor family. The AP-1 transcription factor family plays a role in a variety of processes including angiogenesis, cell proliferation, and cytotrophoblast differentiation/fusion. Carboxypeptidase, vitellogenic–like *(CPVL)* was both an altitude-associated and an ancestry-associated DEG, demonstrating decreased expression at high altitude and increased expression in Andean placentas.

**Table 3:** Fold change of Illumina HT12-v4 gene expression microarray probes that are significantly differentially expressed (FDR < 0.1) associated with altitude.

| Probe ID | Gene Symbol | Fold Change | FDR |
| --- | --- | --- | --- |
| ILMN_1669523 | *FOS* | -2.04 | 0.045 |
| ILMN_1762899 | *EGR1* | -1.9 | 0.055 |
| ILMN_1751607 | *FOSB* | -1.63 | 0.080 |
| ILMN_1806023 | *JUN* | -1.56 | 0.068 |
| ILMN_1892403 | *SNORD13* | -1.47 | 0.068 |
| ILMN_2242463 | *CTSC* | -1.35 | 0.098 |
| ILMN_1778977 | *TYROBP* | -1.31 | 0.084 |
| ILMN_1727532 | *OLFML3* | -1.3 | 0.069 |
| ILMN_1682928 | *CPVL* | -1.3 | 0.075 |
| ILMN_1656011 | *RGS1* | -1.28 | 0.069 |
| ILMN_1682717 | *IER3* | -1.27 | 0.075 |
| ILMN_1812073 | *ATP6V1B1* | -1.25 | 0.075 |
| ILMN_2075927 | *STK40* | 1.26 | 0.068 |
| ILMN_1727589 | *SULT2B1* | 1.29 | 0.055 |
| ILMN_1763749 | *GUCA2A* | 1.29 | 0.075 |
| ILMN_1689817 | *LCOR* | 1.29 | 0.098 |
| ILMN_1772809 | *SLC4A1* | 1.29 | 0.055 |
| ILMN_1677684 | *BTBD16* | 1.3 | 0.080 |
| ILMN_1713751 | *ADAM19* | 1.3 | 0.069 |
| ILMN_2205963 | *C10orf54* | 1.31 | 0.068 |
| ILMN_1784333 | *SECISBP2L* | 1.32 | 0.055 |
| ILMN_2376502 | *RHOBTB1* | 1.32 | 0.070 |
| ILMN_1765332 | *TIMM10* | 1.33 | 0.069 |
| ILMN_1695187 | *GYPE* | 1.34 | 0.068 |
| ILMN_1746578 | *SLC23A2* | 1.35 | 0.055 |
| ILMN_1717262 | *PROCR* | 1.35 | 0.069 |
| ILMN_1695157 | *CA4* | 1.37 | 0.069 |
| ILMN_1810420 | *DYSF* | 1.38 | 0.001 |
| ILMN_1740604 | *RAB11FIP5* | 1.4 | 0.069 |
| ILMN_1763666 | *ALDH3B2* | 1.42 | 0.067 |
| ILMN_1759155 | *IFIT1B* | 1.42 | 0.068 |
| ILMN_1707734 | *NNAT* | 1.43 | 0.075 |
| ILMN_1704376 | *GLDN* | 1.44 | 0.075 |
| ILMN_1696512 | *AHSP* | 1.52 | 0.068 |
| ILMN_1713266 | *FAM46C* | 1.57 | 0.068 |
| ILMN_1773006 | *FABP4* | 1.67 | 0.068 |

**Figure 1:**
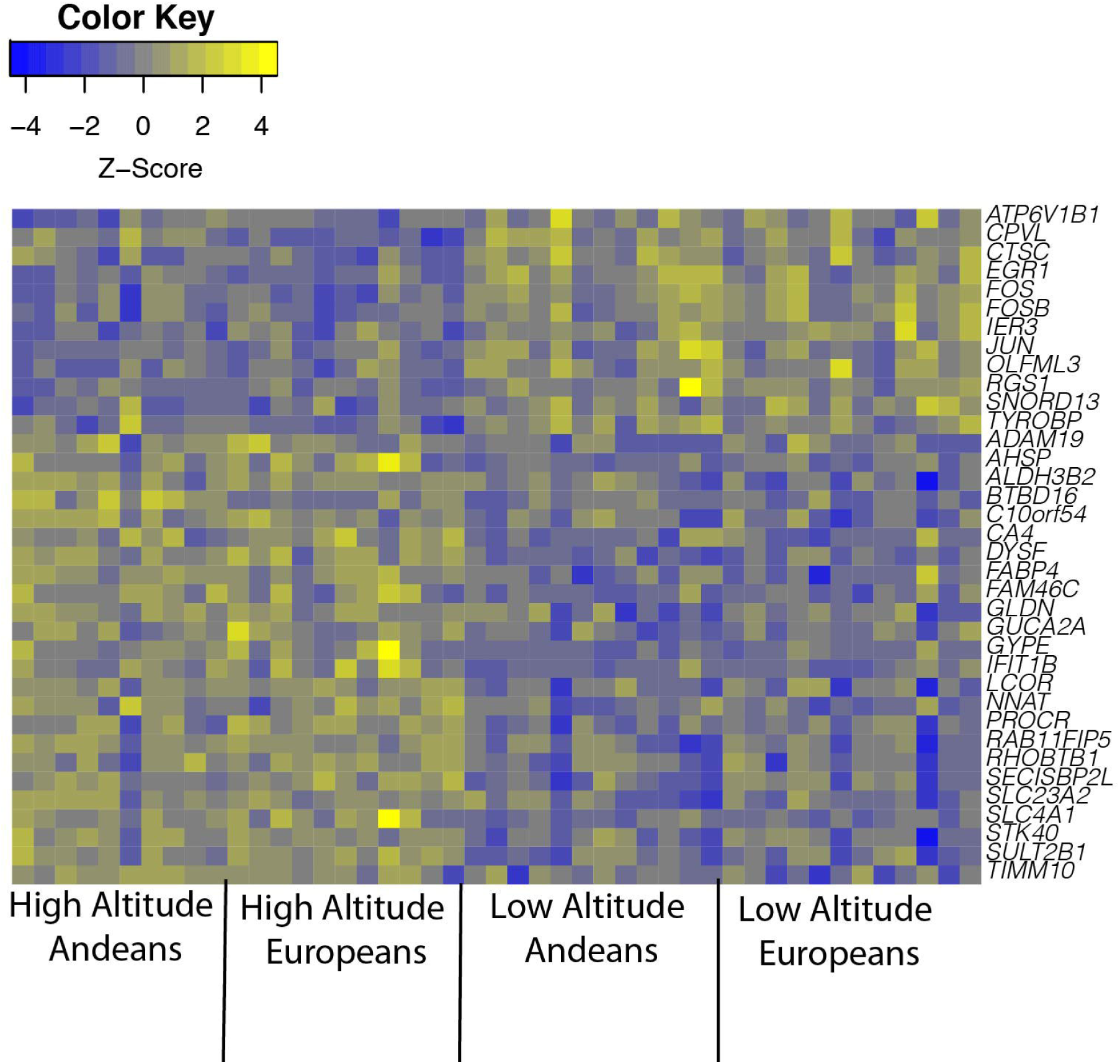
Heat map of Z-scores for each altitude-associated differentially expressed genes. Z-scores calculated from gene expression of each altitude-associated differentially expressed gene. The blue pixels represent negative z-scores (i.e. expression level for that sample is less than the mean expression) and the yellow pixels represent samples with positive z-scores (i.e. expression levels for that sample is greater than the mean expression level). A gene was considered to be differentially expressed with a FDR<0.1 and an absolute fold change≥1.25 between individuals residing at high and low altitude.

To test for enriched KEGG pathways amongst the DEGs, over-representation analysis was conducted using GOstats [26]. Enriched pathways (FDR<0.05) among altitude-associated DEGs include osteoclast differentiation (KEGGID: hsa04380) and rheumatoid arthritis (KEGGID: hsa05323); each involves DEGs within the activator protein 1 (AP-1) transcription factor family including *JUN, FOS, FOSB*, and early growth response 1 *(EGR1)*. Enriched pathways among ancestry-associated DEGs included rheumatoid arthritis (KEGGID: hsa05323), allograft rejection (KEGGID: hsa05330), and graft-versus-host disease (KEGGID: hsa05332). These pathways include DEGs that are involved in inflammation and angiogenesis including *HLA-A, HLA-DRB1, CCL2*, and *CCL3L3*.

### 3.3. Correlation of DNA Methylation with Gene Expression

1,065 Illumina HM450k methylation microarray probes, each designed to measure methylation at a specific CpG site, were located within the promoter or gene body region of the 69 ancestry- and altitude-associated DEGs. 121 methylation sites correlated significantly with gene expression levels when corrected for multiple hypothesis testing (FDR<0.1) (Supplementary Table 3). 106 methylation sites had a significant correlation with expression levels of 17 ancestry-associated DEGs while 15 methylation sites correlated significantly with expression levels of 8 altitude-associated DEGs (Supplementary Table 3, Supplementary Figure 2). Seventy-two percent (28/39) of the methylation probes located in a promoter region had a negative correlation with gene expression. Among these 39 probes, eighty-nine percent (8/9) of the probes among altitude associated DEGs had a negative correlation while sixty-six percent (20/30) of the probes among ancestry-associated DEGs had the same negative correlation. A negative correlation suggests that higher methylation values were associated with lower gene expression and vice versa. Some ancestry-associated DEGs that had a significant negative correlation between gene expression and promoter DNA methylation included the cell surface markers *HLA-DRB1, HLA-A, CPVL*, and *LYVE1*. Altitude-associated DEGs showing a negative correlation included genes involved in the response to oxidative stress and inflammation such as gliomedin *(GLDN)*, TYRO protein tyrosine kinase binding protein *(TYROBP)*, dysferlin *(DYSF)*, and *CPVL*.

In addition to promoter region methylation, 69% (58/84) of the methylation probes that correlated significantly within gene bodies showed a pattern of negative correlation with gene expression. We also calculated correlation coefficients within CpG Islands located within gene bodies (i.e. >200 bp sequence within a gene body with > 60% CpGs). Ninety-seven percent (38/39) CpG sites within CpG islands located within gene bodies showed a negative correlation with gene expression. There were more significant correlations between gene expression and DNA methylation among ancestry-associated DEGs than altitude-associated DEGs (Fisher’s exact p=0.0242, Table 4).

**Table 4:** 2 × 2 contingency table of genes that may be regulated by DNA methylation or not regulated by DNA methylation among altitude and ancestry associated differentially expressed genes.

|  | **Ancestry** | **Altitude** |
| --- | --- | --- |
| **Regulated by DNA methylation** | 17 | 8 |
| **Not Regulated by DNA methylation** | 17 | 28 |

## 4. Discussion

We present here a comprehensive analysis of mRNA expression and DNA methylation in villous placental tissue from 45 Bolivian individuals of primarily Andean or European ancestry from high versus low altitude. We have shown that (1) a total of 69 genes were differentially expressed in the placenta in association with either altitude or ancestry, and (2) that epigenetic regulation of gene expression was more commonly associated with ancestry (Andean vs. European) as opposed to the environmental stressor of interest (altitude); and (3) how the DEGs might relate to the structural differences observed in the placentas themselves, which confer fetal growth advantages of Andean relative to European high-altitude inhabitants.

### 4.1. Ancestry-associated Differentially Expressed Genes

Of the 34 ancestry-associated DEGs, 24 were more highly expressed in Andean compared to European placentas (Table 2, Figure 2, Supplementary Table 1). Among these ancestry-associated DEGs are several genes associated with inflammation including two inflammatory chemokines, c-c motif chemokine ligand 2 *(CCL2)* and c-c motif chemokine ligand 3 like 3 (CCL3L3). The ancestry-associated increase in expression of inflammatory chemokine ligands, *CCL2* and *CCL3L3*, would be expected to attract increased numbers of immune cells to the placenta (e.g. natural killer cells, macrophages, and monocytes), important in normal invasion and remodelling of the maternal-fetal interface [31]. Previous work has shown increased pro-inflammatory cytokines in the circulation of pregnant women at high altitude [32].

**Figure 2:**
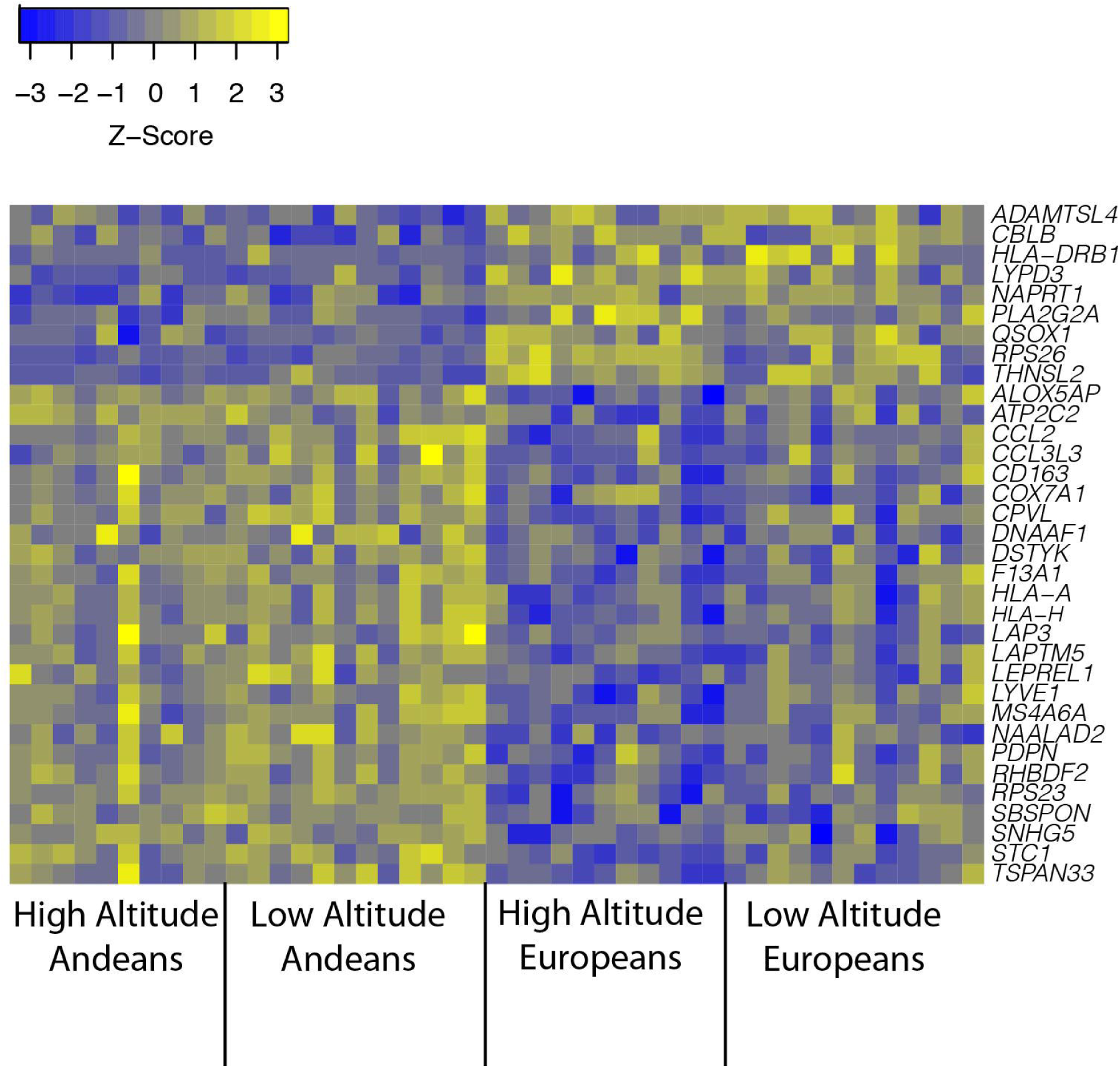
Heap map of Z-scores for each ancestry-associated differentially expressed genes. Z-scores calculated from gene expression of each ancestry-associated differentially expressed gene. The blue pixels represent negative z-scores (I.e. expression level for that sample is less than the mean expression) and the yellow pixels represent samples with positive z-scores (i.e. expression levels for that sample is greater than the mean expression level). A gene was considered to be differentially expressed with a FDR<0.1 and an absolute fold change≥1.25 between individuals of Andean and European descent.

In addition to these inflammatory genes, several markers of fetally derived pro-angiogenic macrophages (Hofbauer cells), including CD163 molecule *(CD163)*, lymphatic vessel endothelial hyaluronan receptor 1 *(LYVE1)*, and major histocompatibility complex class 1 A (*HLA-A)*, displayed higher expression levels in Andeans compared to Europeans. Hofbauer cells, i.e. placental macrophages, can secrete pro-angiogenic factors which may contribute to the enhanced vascularity of the high-altitude Andean placenta and contribute to enhanced nutrient and gas exchange [33]. We have shown that the placenta engages in mitochondrial hypometabolism under high altitude conditions, and regulated by HIF-1 alpha [34, 35]. HIF-1 alpha regulates expression of chemokines, including *CCL2* and *CCL3L3* which may result in increased angiogenesis [36]. We therefore speculate that the genes we have identified related to the innate immune response may be permissive for the greater angiogenesis that we and others have observed in Andean placentas at high altitude [11,13].

Andean women had a robust, altitude-associated increase in placental capillary surface area, length and volume, despite general simlarity in placental weight and volume aross the 4 groups. Increases in these functional parameters were 19-71% greater in Andean than European placentas for the same vascular variables. Functionally, one would expect greater capillary surface area and length to be associated with increased maternal-fetal nutrient transport and gas diffusion, whilst volume might be expected to relate to blood flow. We have no direct measures of diffusion, although extensive analysis of fetal blood gas parameters and oxygen delivery suggested the Andean fetus is advantaged relative to the European migrants [8]. At the same time, and in these same pregnancies, we showed greater umbilical venous blood flow in the Andean vs. European fetus at high altitude, which may be related to their greater capillary volume [8]. These data indicate a pronounced Andean advantage in the placental structural response to the chronic hypoxia of high altitude, centered on angiogenesis, and consistent with many of the ancestry-associated DEGs

### 4.2. Altitude-associated Differentially Expressed Genes

Of the 36 altitude-associated DEGs, 10 DEGs had lower expression levels at high altitude compared to low altitude (Table 3). Three of those genes, fos proto-oncogene *(FOS)*, fosb proto-oncogene *(FOSB)*, and jun proto-oncogene *(JUN)* had the greatest negative fold-change in high altitude placentas. These genes are part of the activator protein 1 (AP-1) transcription factor complex. The AP-1 complex is a family of transcription factors formed by the dimerization of products from the *FOS* and *JUN* gene families [37]. AP-1 transcription factors regulate the expression of genes involved in cell proliferation, cell death, and cellular transformation [38]. This is consistent with molecular analyses addressing regulation of the cell cycle and apoptosis under in vivo (preeclampsia, IUGR, altitude) and in vitro low oxygen conditions [39-42]. Previous studies have found an increase in cytotrophoblasts in high altitude placenta [43, 44], interpreted as hypoxia-induced proliferation. However, molecular data supports the excess cytotrophoblast cells are there due to a reduced rate of cytotrophoblast fusion into the syncytium. In essence syncytialization is delayed or slowed and apoptotic stimuli are diminished in the high altitude placenta.

Decreased expression levels of genes forming the AP-1 transcription factor complex indicate that there is less AP-1 transcription factor available to bind to its target genes. The reduction in expression levels of four AP-1 target genes at high altitude supports less availability of the AP-1 transcription factor. These four genes include early growth response 1 *(EGR1)*, olfactomedin like 3 *(OLFML3)*, regulator of G protein signaling *(RGS1)* and immediate early response 3 *(IER3)* all of which have been shown to be involved in cell proliferation, suggesting that decreased expression may result in reduced cell proliferation [45-48]. The reduced cell proliferation may also contribute to reduced cytotrophoblast fusion into the syncytium [49, 50].

Prior studies of gene expression in Tibetans and Andeans implicate the HIF pathway as a critical target of natural selection, and hence these pathways were examined carefully in this study [51, 52]. The altitude associated DEGs include genes involved in the response to hypoxia and the induction of the hypoxia-inducible factor (HIF) pathway. DEGs associated with hypoxia and the HIF pathway show higher gene expression among residents of high vs. low altitude. The DEGs associated or induced by HIF induction with higher expression at high altitude include family with sequence similarity 46 member C *(FAM46C)*, solute carrier family 4 member 1 (Diego blood group) *(SLC4A1)*, and alpha hemoglobin stabilizing protein *(AHSP)* (Figure 1, Table 3) [53-55]. We have previously shown increased HIF-1 alpha gene expression, protein levels, and increases in the products of HIF’s target genes in placentas of non-indigenous peoples residing at high altitude [56, 57]. Placental gene expression for HIF-1 alpha *(HIF1A)*, HIF-2 alpha *(EPAS1)*, and HIF-3 alpha *(HIF3A)* in this study demonstrated low total variance (σ^2^ < 0.1186) and therefore were not subject to genome wide hypothesis testing. There was however greater expression of the HIF-3 alpha gene product among Andeans compared to Europeans regardless of altitude (p=0.01). We also found that there was greater HIF-1 alpha gene expression among Europeans vs. Andeans residing at high altitude (interaction p-value = 0.02). It is unlikely that the gene’s nominal p-value would hold up to the multiple hypothesis testing required in genome wide hypothesis testing, but this observation illustrates an interesting point. Several of the HIF-3alpha isoforms can attenuate HIF-1alpha binding to HREs in target genes [58, 59] and thereby modulate HIF-1alpha-dependent gene transcription. Studies of gene expression from leukocytes of Tibetans and Andeans indicate natural selection has favored variants of genes in the HIF-pathway that would be expected to lower the Po^2^ at which a HIF response is stimulated. This appears counter-intuitive unless one posits that this lowered threshold for stimulation of a HIF response contributes to maintenance of sea-level values for some physiological parameters, despite the high altitude environment, which is as good a measure of adaptation as any other [51, 52]. That HIF-3alpha can attenutate HIF-1alpha target gene transcription and is elevated in placentas of Andeans regardless of altitude suggests it may act to further ‘fine-tune’ HIF-1alpha responses. Given that vascularity in low altitude Eurpeans and Andeans is similar, these differences in HIF-related gene expression at high altitude begs the question of whether the increased HIF-3alpha and relatively lower, but still elevated, HIF-1alpha levels plays a role in the Andean advantage in placental vascular response to high altitude hypoxia [59].

### 4.3. Correlation between Genes that were Differentially Expressed Associated with Altitude or Ancestry and Methylation Patterns

There were more ancestry-associated DEGs (n=17) that correlated with DNA methylation than altitude-associated DEGs (n=8, Table 4, Supplementary Table 3). This significant difference (Fisher’s exact p = 0.0242) supports our hypothesis that methylation plays an important role in placental adaptation to high altitude. DNA methylation changes can be introduced within a population at a faster rate than gene expression differences due to evolutionary selection for extant or new sequence polymorphisms [60]. Placental DNA methylation would be favored over sequence variation as a readily modifiable way to alter gene expression without inducing a permanent genetic change in the germ cell line. This has been reconized in the obstetric literature [61]. It is one additional way in which phenotypic plasticity, which is amply well documented among Andeans in quantifiable traits such as chest dimensions and lung volumes, is favored over genetic changes that would be permanent.

It is well known that hypermethylation in the promoter region of genes typically results in lower gene expression and an inverse correlation between DNA methylation and gene expression [62]. The inverse correlation is likely due to DNA methylation limiting the binding of transcription factors and transcriptional complexes to their respective binding sites [63]. We have shown that among our DEGs, 72% (28/39) of methylation probes located within promoter regions had methylation values that were inversely correlated with gene expression with the remaining 28% (11/39) of promoter methylation sites having a positive correlation. We also found negative correlations between gene expression levels and DNA methylation within 97% (37/38) of the methylation probes located within CpG islands of gene bodies. These findings suggest that these CpG islands may be acting as alternative promoters since CpG islands are typically located within promoter regions [64, 65]. Transcription at alternative promoters may lead to isoforms that are unique to either an ancestral group or individuals exposed to an environment such as high altitude [66, 67]. The use of intragentic CpG islands as tissue specific alternative promoters has been among cells that are part of the hematopoetic steam cell lineages as different cell types have different methylation patterns within CpG islands [67].

In contrast to DNA methylation in promoters and CpG islands within gene bodies (i.e. introns and exons), increased DNA methylation within non-CpG island gene bodies is increasingly becoming associated with increased mRNA abundance and a positive correlation between gene expression and DNA methylation [68]. However, our results show only 57% (25/43) of the methylation sites within the gene body, excluding those within CpG islands, had methylation values that correlated positively with gene expression. This suggests that 43% of non-CpG island gene body DNA methylation may be acting in a similar fashion to promoter region methylation [62]. The genes we found that had decreased promoter methylation and consequently higher gene expression among Andeans, are associated with inflammation and pro-angiogenic macrophages. Those ancestry-associated DEGs with decreased promoter methylation include carboxypeptidase, vitellogenic-like *(CPVL)*, lymphatic vessel endothelial hyaluronan receptor 1 *(LYVE1)*, prolyl 3-hydroxylase 2 *(LEPREL1)* and N-acetylated alpha-linked acidic dipeptidase 2 *(NAALAD2)* [69-73]. This may contribute to the increased angiogenesis in Andeans compared with Europeans placentas at high altitude. Gene expression differences associated with altitude may be less dependent on epigenetic mechanisms and more dependent on the availability of transcription factors (e.g. AP-1 binding), or sequence variation favored by natural selection. Seven altitude-associated DEGs had expression levels that correlated significantly with DNA methylation including genes involved in cellular and membrane repair such as gliomedin *(GLDN)* and dysferlin *(DYSF)*.

This is the first study, to our knowledge, that investigates how placental gene expression differs among distinct populations of different biogeographic ancestry, which gene expression differences may be regulated by DNA methylation, and how these mechanisms may differ in the setting of reduced oxygen tension. It is also the first to consider these differences in relation to structural/functional dfferences in the target organ. It differs from prior studies that have identified polymorphisms found in DNA from leukocytes, which are likely unrepresentative of tissue or cell-specific gene expression patterns [52, 74]. The gene variants identified in those studies have not yet been examined in a functional context nor have pre- or post-translational modifications to the proteins encoded by the genes of interest been tested. Moreover, DEGs based on functional tissues of impact on reproductive outcomes are likely not reflcted in DEGs derived from hemtopoietic cells, which have a biological turnover of 3 months, and are more likely to reflect short-term, rather than long term, reproductively important outcomes. The study reported here provides insight into how DNA methylation may be regulating gene expression in the placental tissue of Andeans, which, unlike Tibetans, may have significant exposure to low altitude environments through culture and commerce. It suggests that Andean placentas may be adapting to the high altitude environment, seen by a preservation in birth weight, through epigenetic modifications as opposed to sequence variation.

A major limitation of this study is the small sample size, which restricts the power of our analysis by limiting the number of probes that would be significant after adjusting for multiple hypothesis testing. To account for this, we focused our analysis on only the top 5% most variable probes. Recent literature on altitude-associated genetic variation revealed polymorphisms in genes related to the hypoxia-inducible factor nuclear transcription family of genes (i.e. *HIF1A, EPAS1, HIF3A)* between high and low altitude populations [52, 75]. The fact that we did not see differential expression in the same genes in which leukocyte DNA polymorphisms were identified may be associated with the fact that the placenta of adapted individuals at high altitude receives the same amount of oxygen as the low altitude placenta, albeit at markedly lowered blood flow [8]. Other limitations include the range of RIN scores used in this study which may affect the ability of the respective cDNA to hybridize to the microarray chip affecting expression levels. Studies asking whether there has been natural selection of genetic polymorphisms or novel sequence variants in high altitude populations are limited to leukocytes. In our study, HIF-related genes did not achieve the level of statistical significance required for further exploration. Larger sample sizes may remedy this. Finally, ancestry differences may have affected hybridization, as the microarray chips were designed using primarily European sequences.

## Conclusion

We have identified differences in gene expression within the term human placenta in response to two different factors, altitude (high vs. low) and biogeographic ancestry (Andean vs. European). We have attempted to link the gene expression changes to differences in placental structure and consequently birth weights at high altitude. The approach is warranted because of the strong correlation between placental size and structure with pregnancy outcomes such as birth weight, preeclampsia and IUGR [76-79]. We found that the ancestry-related expression differences are related to inflammatory processes and angiogenesis. By contrast, the altitude-related expression differences are more-closely linked to changes in the AP-1 transcription network. These may be linked to decreased cytotrophoblast turnover (and/or impaired syncytialization).

Lastly, we have shown that DNA methylation appears to regulate the ancestry-associated DEGs more than the altitude-associated DEGs. Taken together, the results suggest that the phenotypic plasticity conferred by methylation over sequence variation in genetic polymorphisms is a preferred evolutionary strategy. Epigenetic changes, because they can be environmentally induced and presumably absent or reversed in an alternative environment, permit greater flexibility in the face of stressful/changeable environments. These data also support that organ-specific or target tissue evaluation of gene expression is required in the assessment of genetic adaptation to environmental stress.

## Supplement List

**Supplementary Table 1:** Tables of differentially expressed genes associated with altitude and/or ancestry

**Supplementary Table 2:** Validation of gene expression microarray by qRT-PCR

**Supplementary Table 3:** Results of the Pearson’s correlation test between DNA methylation and gene expression

**Supplementary Figure 1:** Scatterplot of variance of Illumina HT12v4 gene expression microarray probes

**Supplementary Figure 2:** Correlation of gene expression with methylation values of differentially expressed genes

Differentially expressed genes with at least one methylation site that significantly correlates with gene expression values sorted by its location with the right most sites being located on chromosome 1. The green circles represent CpG sites located within the gene body. The orange circle represents CpG sites located within the promoter region. The purple circles represent CpG sites that are located within CpG islands of a differentially expressed gene.

